# Structural Determinants of the Interactions Between Influenza A Virus Matrix Protein M1 and Lipid Membranes

**DOI:** 10.1101/369462

**Authors:** C.T. Höfer, S. Di Lella, I. Dahmani, N. Jungnick, N. Bordag, S. Bobone, Q. Huang, S. Keller, A. Herrmann, S. Chiantia

## Abstract

Influenza A virus is a pathogen responsible for severe seasonal epidemics threatening human and animal populations every year. One of the ten proteins encoded by the viral genome, the matrix protein M1, is abundantly produced in infected cells and plays a structural role in determining the morphology of the virus. During assembly of new viral particles, M1 is recruited to the host cell membrane where it associates with lipids and other viral proteins. The structure of M1 is only partially known. In particular, structural details of M1 interactions with the cellular plasma membrane as well as M1–protein interactions and multimerization have not been clarified, yet.

In this work, we employed a set of complementary experimental and theoretical tools to tackle these issues. Using raster image correlation, surface plasmon resonance and circular dichroism spectroscopies, we quantified membrane association and oligomerization of full-length M1 and of different genetically engineered M1 constructs (i.e., N- and C-terminally truncated constructs and a mutant of the polybasic region, residues 95-105). Furthermore, we report novel information on structural changes in M1 occurring upon binding to membranes. Our experimental results are corroborated by an all-atom model of the full-length M1 protein bound to a negatively charged lipid bilayer.

## INTRODUCTION

Influenza A viruses (IAV) circulate in human and animal populations, posing a significant threat to global health. They are responsible for severe pathologies with high mortality during seasonal epidemics and occasional pandemic outbreaks [1]. IAVs belong to the family of *Orthomyxoviridae* and are negative-sense single-stranded RNA viruses [2]. The segmented viral genome, in form of eight viral ribonucleoprotein complexes (vRNPs), is enclosed by the viral lipid envelope [3]. The matrix layer underneath the viral membrane is formed by the matrix protein M1, which consists of 252 amino acid (aa) residues [4-8]. M1 is the most abundant protein in virus particles, stabilizes the virion structure and is considered to be a determinant of viral morphology [9-13]. Electron micrographs demonstrate tight association of the matrix layer with the viral lipid membrane [5]. M1 is abundantly produced in infected cells and distributes throughout cytoplasm and nuclear compartment. In addition to its role as a structural protein, M1 has other functions during the viral replication cycle. It is involved in nucleocytoplasmic transport of the viral genome and, therefore, plays a role in coordinating nuclear-localized replication and assembly of progeny viruses at the plasma membrane (PM) [14-19]. M1 is furthermore considered to be a key player during virus assembly because of its ability to associate with all other structural components of virus particles, including the genomic vRNP complexes [20-22], the three viral transmembrane proteins (i.e. hemagglutinin (HA), neuraminidase (NA) and the proton channel M2) [23-26] and the lipid membrane itself [5, 21, 27, 28]. Available data suggest a model of virus assembly in which M1 is recruited to the budding site at the PM through binding to negatively charged lipids and interactions with the cytoplasmic tails of HA, NA or M2 [12]. Multimerization of M1 and multiple protein– protein and protein–lipid interactions can then interconnect the various components and define the site of virus formation [25, 29-31]. Bending of the membrane and incorporation of the viral genome finally lead to formation and release of new virus particles [12].

Several biophysical studies on model membranes have demonstrated specific binding between M1 and negatively charged lipids such as phosphatidylserine (PS) [5, 21]. We have previously shown that M1 binds to PS clusters in the PM of infected cells [29] and, furthermore, M1–lipid interaction induces the formation of M1 multimers bound to lipid membranes, both in model and cellular systems [32]. The processes of M1 multimerization and membrane binding, which are both required during virus formation, are apparently connected. Recently, we have proposed a model according to which binding to lipids might induce conformational changes in M1 that trigger or, at least, enhance protein–protein interaction and multimerization [32]. However, the effect of M1–lipid binding on the M1 structure has not been investigated yet. Moreover, it is not clear which residues in M1 mediate the association with lipid membranes and the subsequent M1 multimerization. While crystal structures of the relatively compact M1 N-terminal domain have been solved at pH 4.0 and 7.0, both the structure and the exact orientation of the more flexible and disordered C-terminal domain within the full-length protein remain unclear [33-37]. A recently published crystal structure of the matrix protein from another orthomyxovirus, the infectious salmon anaemia virus (ISAV), provides first clues on the 3D structure of full-length IAV M1 [38].

Since M1 membrane binding and multimerization are two physiologically related processes in the context of virus assembly, they need to be studied in combination to account for the connection between both processes and to allow dissecting the contributions of different structural domains to each of these interactions. Raster image correlation spectroscopy (RICS) is a fluorescence-based approach that can be used to study protein binding to supported lipid bilayers (SLB), thereby providing a powerful tool to investigate M1 multimerization in the context of membrane binding, as we have previously demonstrated [32]. In the present study, we employed RICS to analyze and compare different structural domains of M1 with regard to membrane association and oligomerization: the truncated M1 constructs M1-N (aa 1–164) and M1-C (aa 165–252) as well as the polybasic domain (PBD) mutant M1_m_ (in which all basic residues in positions 95–105 were replaced by alanine residues [39]) were compared to wild-type (wt) M1. Additionally, surface plasmon resonance (SPR) measurements were performed as a complementary approach to determine the affinity of these constructs to negatively charged lipid membranes in the absence of any chemical modifications that might perturb protein–lipid interaction. In order to investigate conformational changes that might occur in M1 and each of its domains upon membrane interaction, we performed circular dichroism (CD) experiments. For validation of our experimental results and further insight into the details of M1–membrane interaction, we conducted molecular dynamics simulations (MDS), yielding a model of M1–membrane interaction as a dynamic process at the atomic level [40, 41]. Through combination of the different approaches, we provide a molecular interpretation of M1–lipid interaction according to which i) M1 stably associates with a negatively charged lipid bilayer via specific residues in its N-terminal domain and ii) protein– lipid interaction and multimerization are connected to changes in protein secondary structure and intra-protein dynamics.

## MATERIALS AND METHODS

### Chemicals

Lipids were purchased from Avanti Polar Lipids (Alabaster, AL, USA) and used without further purification: 1,2-dioleoyl-*sn*-glycero-3-phosphocholine (DOPC), 1,2-dioleoyl-*sn*-glycero-3-phosphoserine (DOPS), 1,2-dioleoyl-triammonium-propane (DOTAP), 1,2-dioleoyl-sn-glycero-3-phosphoserine-N-(7-nitro-2-1,3-benzoxadiazol-4-yl) (N-NBD-DOPS), 1,2-dioleoyl-sn-glycero-3-phosphoethanolamine-N-(7-nitro-2-1,3-benzoxadiazol-4-yl) (N-NBD-DPPE). Restriction enzymes and isopropyl-β-D-thiogalactoside (IPTG) were obtained from Fermentas/Thermo Scientific (Schwerte, Germany). Phusion DNA polymerase was from Finnzymes (Espoo, Finland). Bacto tryptone, Bacto yeast extract and Bacto agar were bought from BD (Heidelberg, Germany). Ampicillin, bovine serum albumin, deoxyribonuclease I (DNase I), dithiothreitol (DTT), imidazole, guanidine hydrochloride, glutathione, lysozyme, *n-*octyl-β-D-glucopyranoside and phenylmethylsulfonyl fluoride (PMSF) were bought from Sigma-Aldrich (Taufkirchen, Germany). Chloramphenicol and ethylenediaminetetraacetic acid (EDTA) were purchased from Serva (Heidelberg, Germany). β-mercaptoethanol and spectroscopy-grade chloroform were from Merck (Darmstadt, Germany). Glucose, salts and sodium/potassium phosphates were acquired from Roth (Karlsruhe, Germany), while Dulbecco’s phosphate buffered saline (DPBS) was acquired from PAN Biotech (Aidenbach, Germany). cOmplete Ultra EDTA-free Protease Inhibitor Cocktail was purchased from Roche (Basel, Switzerland). Alexa Fluor 647 (A647) succinimidyl ester was acquired from Life Technologies (Darmstadt, Germany).

### Plasmid construction for recombinant protein production

The M1 open reading frame (ORF) from influenza/A/FPV/Rostock/34 was amplified from plasmid pHH21-FPV-M (described in [42]) by polymerase chain reaction (PCR) using oligonucleotides M1-*Nde*I-fw (GGGAATTCCATATGAGTCTTCTAACCGAGGTTG–*Nde*I) and M1-*Xho*I-rev (CCGCTCGAGTCACTTGAATCGTTGC–*Xho*I). For amplification of truncated M1 sequences encoding the M1 N-terminus (M1-N, amino acids 1–164) or the M1 C-terminus (M1-C, amino acids 165–252), the PCR primer M1-*Xho*I-rev was replaced by M1-N-rev (CCGCTCGAGTCACTGTCTGTGAGACCGATGC–*Xho*I), or M1-*Nde*I-fw was replaced by M1-C-fw (GGGAATTCCATATGGTGGCTACCACCAATCC–*Nde*I), respectively. The amplified sequences were subcloned into the inducible bacterial expression vector pET15b (Novagen) using restriction endonucleases *Nde*I and *Xho*I, yielding plasmids pET15b-M1,-M1-N and -M1-C. The final plasmid products were then controlled by sequencing. For protein production, chemically competent cells of bacterial strain Rosetta (DE3)pLysS were transformed with these plasmids. Site-directed mutagenesis in order to replace the basic amino acids in the M1 PBD (amino acids 95–105) by alanine residues was performed by two-step overlap-extension PCR. The oligonucleotide GCCGTCAAACTATACGCGGCGTTGGCAGCTGAGATAACATTCTATGG and its reverse complementary oligonucleotide were used in the first step, while CCAAATAACATGGATGCAGCCGTCGCACTATACGCGGCGTTGGCAGC and the reverse complementary sequence were used in the second step. The M1 polybasic mutant (M1_m_) ORF was then inserted into vector pET-15b and transferred into Rosetta (DE3)pLysS for protein production as described above.

### Protein production and purification

M1 protein constructs carrying an N-terminal 6x-His-tag (with a total length of 19 amino acids) were expressed in Rosetta (DE3)pLysS (as described in [32]). Protein production was induced by addition of 0.1–0.4 mM IPTG during exponential growth (OD_600 nm_=0.7) for 3 h at 37 °C. The cells were pelleted and resuspended in ice-cold lysis buffer (16 mM Na_2_HPO_4_, 3 mM KH_2_PO_4_, 500 mM NaCl, 5.4 mM KCl, 200 μg/mL DNase I, 300 μg/mL lysozyme, 5 mM β-mercaptoethanol, 1 mM PMSF, protease inhibitor cocktail, pH 7.4). The lysate was subjected to one freeze-thaw cycle, incubated for 30–60 min at 4°C and sonicated on ice (5 × 20 s). Cell debris was removed by centrifugation at 90,000 g for 20 min. The supernatant was incubated with a TALON metal affinity resin for His-tag purification (Clontech, Saint-Germain-en-Laye, France) for 20–30 min at 4°C under permanent agitation. The suspension was loaded onto a column, and the cleared lysate was allowed to flow through. The column was incubated with equilibration buffer (8 mM Na_2_HPO_4_, 1.5 mM KH_2_PO_4_, 500 mM NaCl, 2.7 mM KCl, 5 mM β-mercaptoethanol, pH 7.4) for another 10 min at 4 °C on a shaker platform. The column was rinsed once more with equilibration buffer, followed by intermediate washing buffer, which was composed of equilibration buffer with 60 mM imidazole, pH 7.2. Protein was finally eluted with equilibration buffer containing 250 mM imidazole (pH 7.2), and eluate fractions were collected. Buffers containing imidazole were always freshly mixed with β-mercaptoethanol followed by pH adjustment. Protein concentrations were determined by absorbance at 280 nm. Proteins were either immediately subjected to fluorescence labeling or stored in elution (10 mM sodium phosphate, 120 mM KCl, 250 mM imidazole, pH 7) buffer at 4°C and used for experiments within two days.

For SPR experiments, the elution buffer containing the protein was exchanged with phosphate buffered saline (PBS: 16 mM Na_2_HPO_4_, 3 mM KH_2_PO_4_, 150 mM NaCl, pH 7.2) by ultrafiltration using Amicon filters (Millipore Ltd., Ireland) with 10 or 3 kDa cut-off.

CD measurements require higher protein concentrations and, therefore, the protein purification protocol was adapted to achieve higher yields by unfolding and refolding of protein constructs from inclusion bodies. Details of the procedure are described in the Supporting Material.

### Fluorescence labeling

Purified M1 constructs were conjugated with the primary amine-reactive dye Alexa Fluor 647 succinimidyl ester. Freshly purified protein (in elution buffer) was incubated with 10-fold molar excess of reactive dye for 18 h at 10 °C, pH 7.2. Free dye was removed by gel filtration with Sephadex G-25. Protein concentration and labeling efficiency were determined by absorbance at 280 nm and 650 nm, respectively. Protein concentrations were between 3 and 20 μM, while labeling efficiencies typically ranged between 0.1 and 0.6 dye molecules per protein.

### Preparation of supported lipid bilayers (SLBs)

Supported lipid bilayers (SLBs) were prepared using the “vesicle fusion method” [43]. For multilamellar vesicle (MLV) formation, 70 mol% DOPC and 30 mol% DOPS were mixed in chloroform and labeled with 0.5 mol% of fluorescent lipid analogue N-NBD-DOPS. The solvent was evaporated, and the lipid film was rehydrated with PBS to a final concentration of 0.7 mg/mL lipid. The MLV suspension was diluted 5-fold and sonicated to form small unilamellar vesicles, 100 μL of which were deposited on a clean glass coverslip within the boundaries of a 7 mm-plastic cylinder that was attached to the glass surface. Vesicle fusion and bilayer formation were induced by addition of 3 mM CaCl_2_. The volume was adjusted to 300 μL and the suspension was incubated for 10 min. Unfused vesicles were removed by addition and removal of 500 μL DPBS, performed 10 times.

### Raster Image Correlation Spectroscopy (RICS)

RICS measurements were performed on an inverted Olympus IX81 microscope equipped with a FluoView FV1000 scanner and confocal detection unit (Olympus, Tokyo, Japan) and coupled to an avalanche photodiode detector (PicoQuant, Berlin, Germany). A 640-nm pulsed diode laser (PicoQuant, Berlin, Germany) was used for excitation in combination with a 60× UPLS Apocromat 1.2 NA water objective and a 650-nm long-pass emission filter.

To analyze M1 binding to the negatively charged lipid bilayers, Alexa Fluor 647-labeled M1 protein constructs were added to the previously prepared supported bilayer to a final concentration of 50 nM (i.e. ~ 1:100 protein/lipid ratio) and incubated for 5 min. Samples were then washed five times by addition and removal of 500 μL of DPBS, immediately followed by recording of RICS data. Data acquisition and analysis was performed as described in Hilsch *et al.* [32]. Further information is also available in the Supporting Material.

### Surface plasmon resonance (SPR)

SPR measurements were performed on a Biacore J (Biacore AB, Uppsala, Sweden) using HPP (XanTec bioanalytics GmbH) or HPA (Biacore) sensor chips. The sensor chip was functionalized with a lipid monolayer according to the protocol provided by the manufacturer. Briefly, the surface of the HPP chip was cleaned using *N*-octyl β-D-glucopyranoside (100 μl, 40 mM in water) for 5 min at a low flow. Lipids were mixed in chloroform at a concentration of 1 mM, with the desired molar ratio (i.e. 70:30 DOPS:DOPC or 100 % DOTAP). The solvent was evaporated under a nitrogen stream and lipids were resuspended by vortexing in PBS (pH 7.4). The dispersion was sonicated to clarity using a bath sonicator (*Emmi* 20 HC, Emag AG, Mörfelden-Walldorf, Germany) for 5 min. The 1 mL 1 mM liposome suspension thus obtained was injected for 60 min. The chip was further washed with a 20 mM NaOH solution multiple times, until producing a stable baseline with signal ranging from 2500 to 3200 response units (RU). This procedure resulted in the formation of a lipid monolayer on the chip surface. The complete coverage of the sensor chip surface by lipids was confirmed by lack of unspecific binding (i.e., <100 RU) of bovine serum albumin (10-min injection of 160 μL of a 1 mg/mL solution in PBS). For all experiments with M1, the control (reference) sensor surface was coated with 100 % DOTAP, for which M1 has very low affinity. The drift in signal for both sample and control flow cells was allowed to stabilize (<5 RU per minute) before any experiments were performed. All solutions were freshly prepared, degassed and filtered through 0.22-μm pores, and measurements were performed at 24 °C in PBS (pH 7.4). Each protein construct was injected at concentrations ranging from 30 nM to ~3 μM (10 μM – 130 μM for M1-C). Protein–lipid association was monitored for 20–23 min (ca. 650 μL volume) to give sufficient time for the association phase to reach near-equilibrium levels. Each sensorgram was then corrected by subtracting the corresponding values recorded for the reference sensor. Measurements were repeated at least as four independent duplicates for each M1 construct, each time using freshly purified protein samples. Binding of each protein construct to DOPS/DOPC monolayers was monitored recording a corresponding sensorgram (i.e. RU as a function of time). Binding curves were, in some cases, characterized by slow association kinetics and complex (i.e. biphasic) behaviour. For this reason, *R_eq_* was simply estimated as the maximum RU recorded after 20 min. *R_eq_* values were then plotted against protein concentrations (*C*) and analyzed with the empirical model described in [44]:

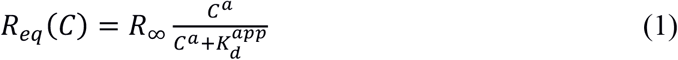

where *R_∞_* is the value of RU at the maximum coverage of bound protein and 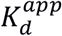 is an apparent dissociation constant of the protein from the lipid monolayer, that is, the protein concentration at which half of the accessible monolayer surface is occupied. Lipid monolayer regeneration was performed after each protein injection to wash away the bound protein by injecting ca. 80 μL of a 10 mM NaOH solution for 1–2 min.

### Preparation of large unilamellar vesicles (LUVs) for circular dichroism (CD) measurements

Lipids at the desired molar ratios (100 % DOPC or DOPC/DOPS (75/25)) and 0.2 mol% N-NBD-DPPE were mixed in chloroform, and the solvent was evaporated under a stream of nitrogen. Lipid films were redissolved in ethanol (one percent of the final volume) and finally resuspended in sodium phosphate buffer (10 mM, pH 7.0) (final lipid concentration 20 mM). Ten freeze–thaw cycles (in liquid nitrogen and 50°C water bath) were performed, and the solution was extruded ten times through polycarbonate membrane filters with a pore size of 0.1 μm (Whatman GmbH, Dassel, Germany).

NBD fluorescence was measured (Aminco-Bowman Series 2 luminescence spectrometer, Thermo Electron Corporation, Germany) to estimate equal amounts of lipid for different preparations. LUVs with a concentration of 2 mM were used within two weeks, 20 mM LUV suspensions were used within three days.

### CD spectroscopy

Protein in sodium phosphate buffer (pH 7.0) was measured at concentrations between 5 and 10 μM in 1-mm cuvettes in a J-720 CD spectrometer (Jasco, Gross-Umstadt, Germany) at 20°C. Independent replicates were also performed in a J-815 CD spectrometer (Jasco). CD spectra were recorded by accumulating five or nine spectra. In the absence of liposomes, CD spectra were recorded between 185 nm and 260 nm. In the presence of liposomes, spectra were acquired between 205 nm and 260 nm, since liposomes gave rise to strong light scattering below 205 nm. The molar ratio between protein and lipid was ca. 1:300. Normalization of the spectra was performed according to the following equation:

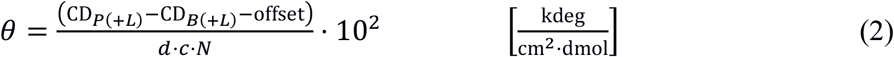

where *θ* is the normalized ellipticity, *CD_P_*_(+*L*)_ the measured ellipticity of the protein (plus liposomes); *CD_B_*_(+*L*)_ the measured ellipticity of the buffer (plus liposomes), *offset* the calculated mean value from 258 nm to 260 nm; *d* the thickness of the cuvette (1 mm; 0.1 mm), *c* the protein concentration in μM, and *N* the number of amino acids residues in the protein construct. UV spectra between 200 and 340 nm were measured prior to CD acquisition to determine protein concentrations according to [45].

### Statistical tests

Statistical significances of differences among data sets were determined using a two-sided t-test with distinct variances (ttest2 routine, Matlab). In the case of asymmetric/skewed data distributions, the logarithms of the corresponding data sets were compared instead. Also, we verified that the use of Wilcoxon–Mann–Whitney test (rank-sum routine, Matlab) on skewed data sets provided similar results.

### Structural prediction and molecular dynamics simulation (MDS) of full-length M1

A detailed description of the methods used to perform MDS can be found in the Supporting Material.

## RESULTS

### The N-terminal domain and PBD are necessary for M1–lipid interaction

In order to dissect the contributions of different M1 domains to the interaction with negatively charged lipid membranes, four different recombinant M1 constructs were produced and purified for use in *in vitro* membrane-binding assays: full-length wild-type (wt) M1, the truncated M1 N-terminal domain (M1-N, aa 1–164), the truncated M1 C-terminal domain (M1-C, aa 165–252) and a full-length M1 mutant (M1_m_) in which all six basic residues of the polybasic domain (PBD, aa 95–105) were replaced by alanine residues.

#### Fluorescence-based quantification of protein binding

In a first approach to study M1– membrane interaction, we used fluorescence laser scanning microscopy to quantify protein binding to well-defined models of the inner leaflet of the plasma membranes (i.e. supported lipid bilayers, SLBs). We have previously shown that this approach can provide quantitative information regarding the amount of bound M1 protein as a function of protein concentration or as a function of negatively charged PS within the membrane [32].

Purified M1 constructs were covalently labeled with the fluorescent dye Alexa Fluor 647 (A647). Labeling efficiencies were typically between 0.1 and 0.6 dye molecules per protein. SLBs were produced with a composition of 70 mol% DOPC and 30 mol% DOPS as a negatively charged lipid. The purified labeled protein was allowed to bind to the membrane for 5 min before unbound protein was washed away. Signal intensities of bound protein were measured by acquiring and averaging several confocal fluorescence images as described in the Materials and Methods section. Fig. 1A shows the average frame of a representative confocal image stack obtained for M1 wt bound to a SLB. Analogous images were obtained for each protein construct (data not shown). Signal intensities of truncated and mutant M1 constructs were then normalized relative to wt M1, as shown in Fig. 1B. Both truncated M1 constructs, M1-N and M1-C, as well as the polybasic mutant M1_m_ associated to significantly lower degree with the negatively charged lipid membrane, compared to M1 wt. These results suggest a concurrent involvement of both N- and C-terminus in membrane binding and a significant role of the PBD in M1 binding to negatively charged lipid membranes. Folding of truncated and mutant M1 secondary structure was verified by circular dichroism (CD) measurements as described below (Fig. 3).

**FIGURE 1.**
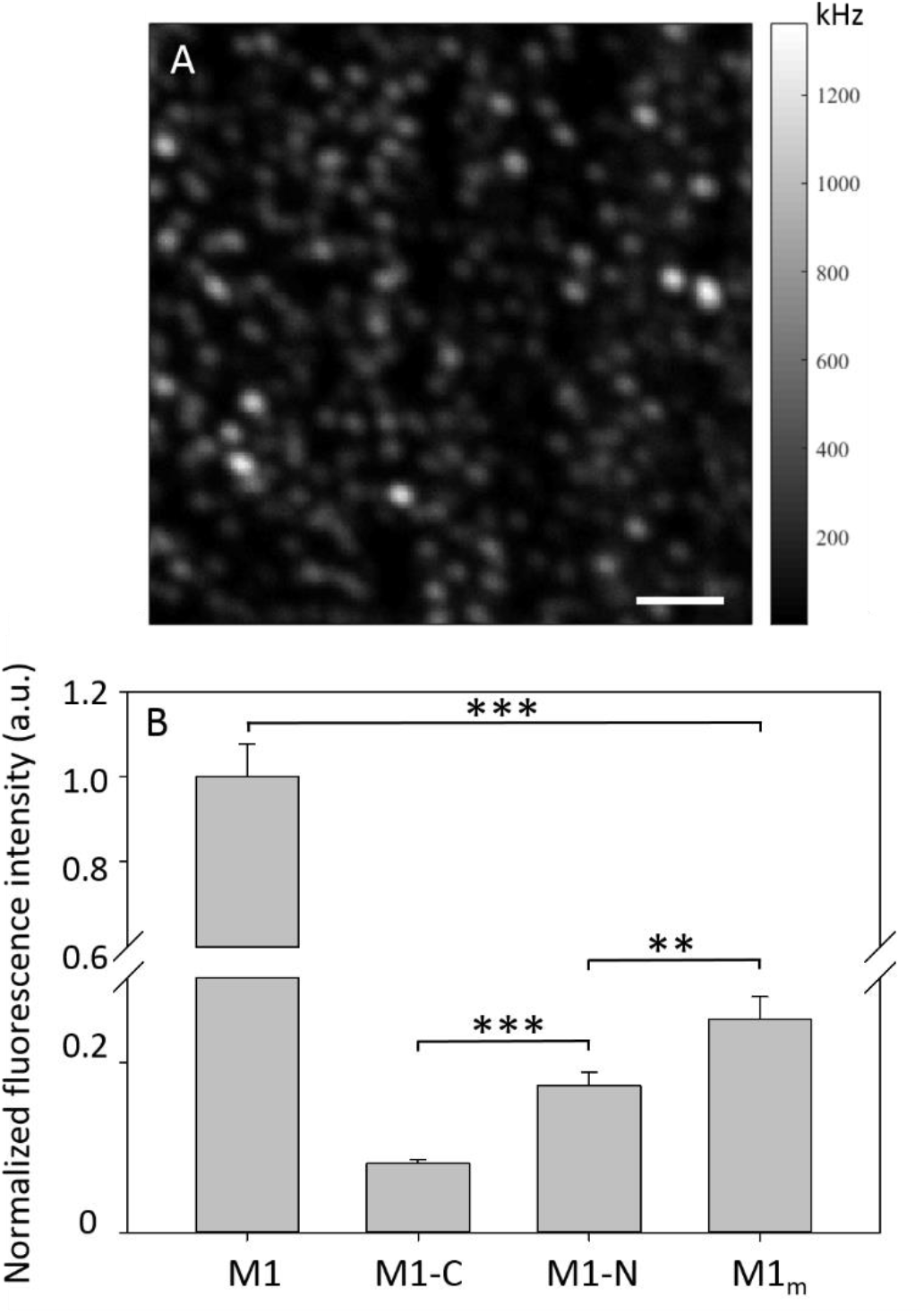
RICS analysis of the membrane binding ability of M1 domains. A: Representative confocal microscopy image of fluorescent M1 wt bound to a PS-containing SLB. The grey-scale indicates the average intensity in each pixel per second. Scale bar is 2 μm. B: Normalized average fluorescence intensity was measured for several confocal stacks of different fluorescently labeled A647-M1 constructs (50 nM) bound to SLBs that contained 70 mol% DOPC and 30 mol% DOPS. Error bars represent the standard error of the mean (n≈100–200 distinct image stacks from 3 (M1, M1-N), 4 (M1-C) or 2 (M1m) independent sample preparations). Asterisks indicate statistically significant differences determined with a two-sided t-test with distinct variances: (^***^ corresponds to p<0.01; ^**^ corresponds to p<0.05). All measurements were conducted at room temperature.

#### SPR-based quantification of protein binding

Binding between lipids and the different M1-based constructs was further characterized using surface plasmon resonance (SPR). This technique has been used in the past to quantify the interactions between lipid membranes and the matrix proteins of several viruses, including HIV and Ebola [44, 46]. The main advantage of this approach is that protein affinity towards model membranes can be quantitatively determined without any need for labeling with, for example, a fluorescent marker. We functionalized the surface of the SPR sensor chip with a monolayer composed of DOPC and DOPS. In order to maximize binding and increase the S/N ratio for constructs with lower membrane affinity (e.g. M1-C, see below), we increased the DOPS nominal concentration to 70 mol%. It is worth noting that, due to the hydrophobic nature of the interaction between sensor chip and lipids, the final relative amount of DOPS in the monolayer might be lower than 70 mol%. In the context of a comparison among different protein constructs, the specific monolayer composition is not expected to play an important role as long as it remains constant among different experiments.

A typical SPR measurement consisted in monitoring the binding of the protein of interest to the lipid monolayer, at increasing concentrations in the range between 3 nM and ~3 μM (10 μM–130 μM for M1-C, due to its low affinity to membranes). Binding of the protein to the monolayer was recorded in a sensorgram as an increase in response units (RU) as a function of time. Fig. 2A shows an exemplary sensorgram obtained for 1 μM M1 wt. All sensorgrams were corrected using a reference sample (see Materials and Methods) and the time point 0 corresponds to the beginning of the injection of the protein solution. The injection/binding phase for each concentration *C* lasted ca. 20 min (see arrows in Fig. 2A), after which a dissociation phase and the regeneration of the monolayer followed (not shown). We focused our attention exclusively on the estimated near-equilibrium value reached after a 20-min binding phase *R_eq_*(*C*). Fig. 2B shows typical binding curves (i.e. *R_eq_ vs. C*) for the different M1 constructs.

**FIGURE 2.**
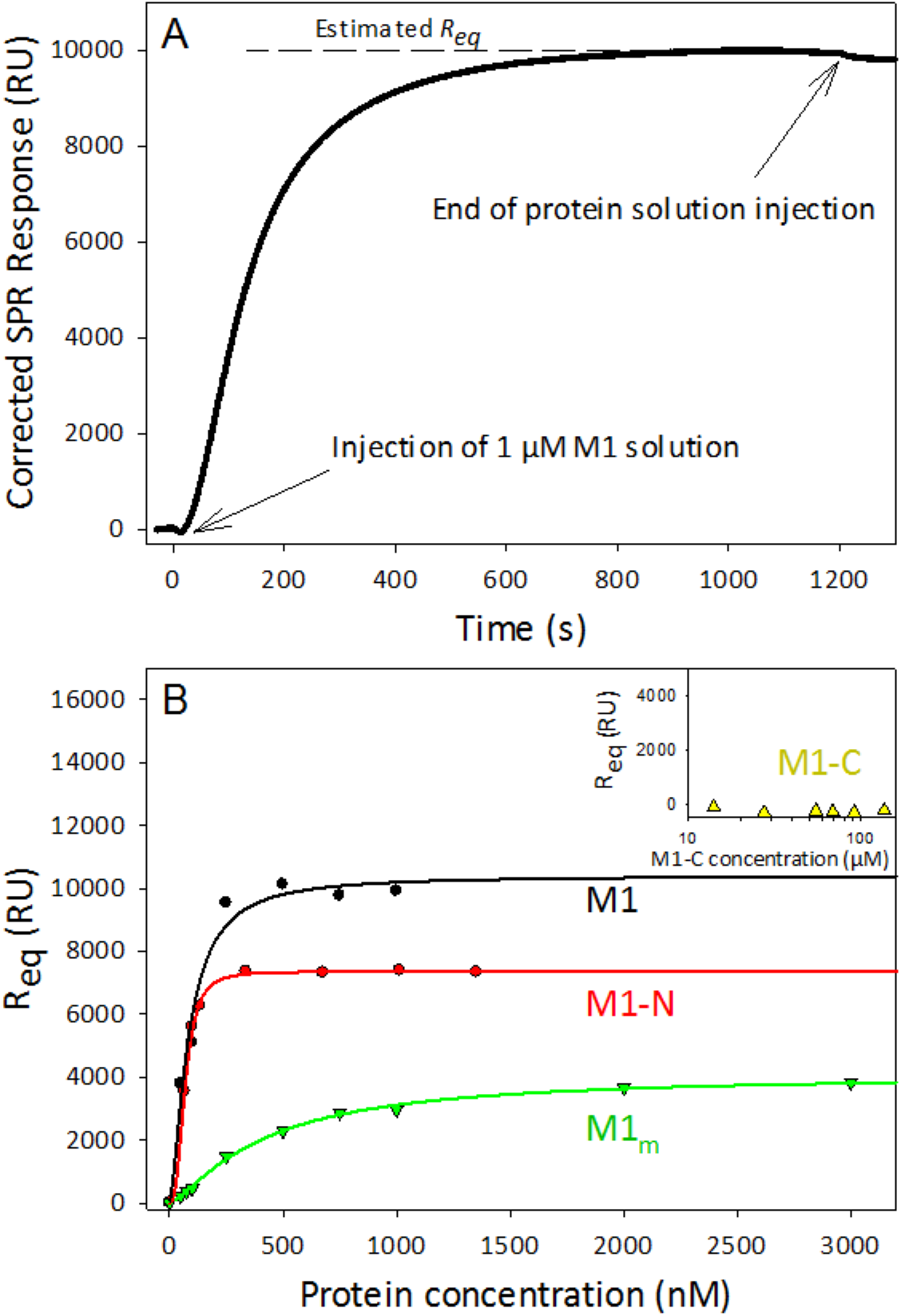
Quantification of M1 binding to DOPC/DOPS membranes via SPR measurements. A: Typical SPR sensorgram obtained for the binding of M1 wt (e.g., 1 μM) to a DOPC/DOPS 30:70 monolayer. The arrows indicate the beginning and the end of protein injection. The SPR response was corrected by subtraction of the reference sample signal. *R*_eq_ was estimated as the value of the SPR response reached at the end of the protein solution injection (i.e. after 20 min). B: Typical binding curves obtained for the different M1 constructs by monitoring *R_eq_* as a function of protein concentration. M1, M1-N and M1_m_ were analyzed in the concentration range 3 nM–3 μM. M1-C concentrations (see inset) were in the 10–100 μM range. The solid lines represent the fitting of the binding curves using an empirical binding model (see text). For each M1 construct, we measured 3– 5 independent protein preparations and analyzed a corresponding number of binding curves. The obtained parameters and the standard deviations are reported in Table 1. All measurements were conducted at room temperature.

**FIGURE 3.**
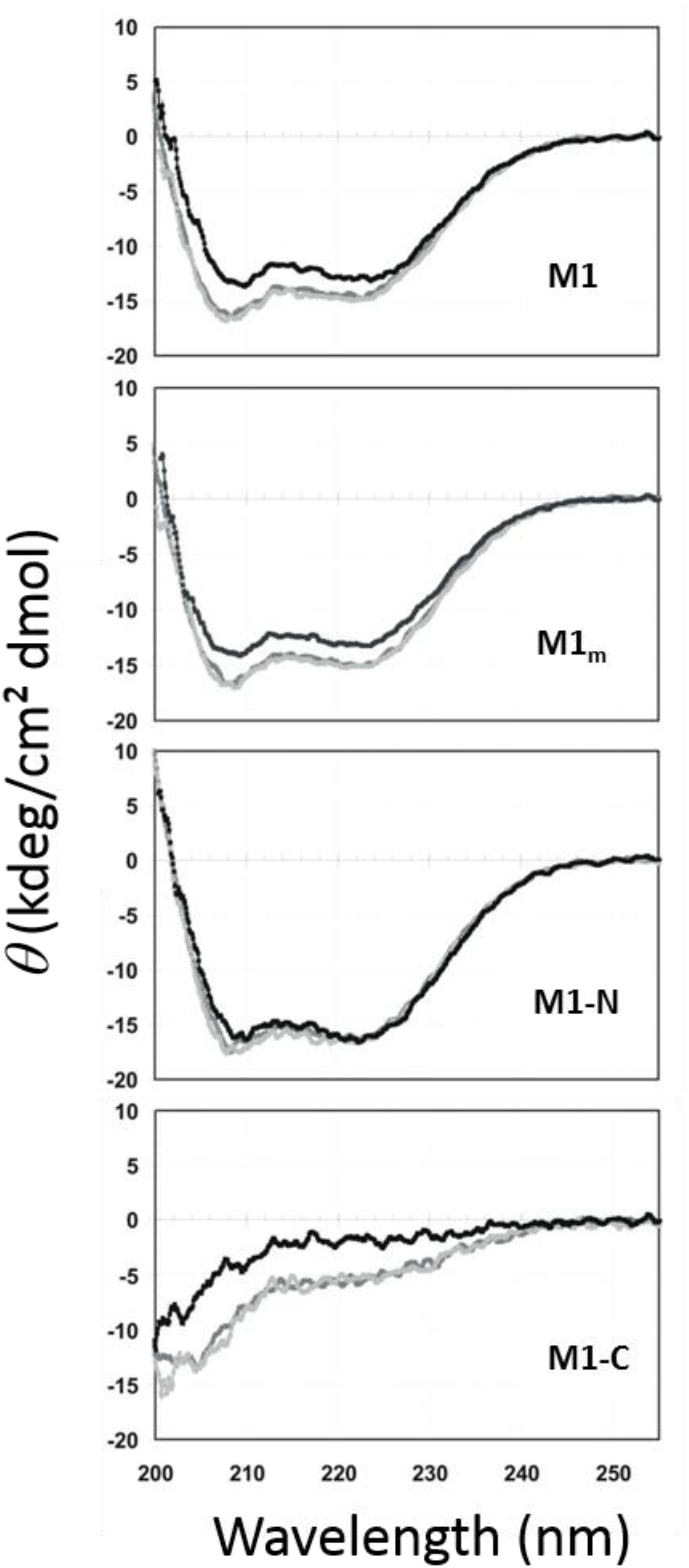
CD spectra of M1 constructs in the presence of lipids. CD spectra were measured for all M1 constructs in buffer (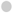) as well as in the presence of DOPC-LUVs (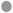) or DOPC/DOPS (7.5/2.5)-LUVs (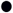). Purified proteins (5-7 μM) were mixed with LUVs (1.6 mM) and incubated for 1 h at room temperature in PBS. Nine spectra were acquired and averaged for each sample and for at least two independently purified protein batches. All measurements were conducted at room temperature.

Similarly to what has been previously reported (44), such curves could not be fitted by a simple Langmuir adsorption isotherm, most likely because of collective adsorption, electrostatic surface effects and protein–protein interaction (i.e. multimerization) during membrane binding. Fitting an empirical binding model (Equation (1)) to the data, as suggested in (44), we were able to analyze the binding of M1, M1-C, M1-N and M1_m_ to DOPC/DOPS monolayers, thus obtaining a maximum protein load (*R*_∞_) and an apparent dissociation constant 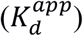. Table 1 summarizes the results thus obtained. A direct comparison of *R*_∞_ (the total amount of bound protein at excess concentrations) can be performed only between constructs with similar mass, that is, M1 wt and M1_m_, and indicates a ca. 3-fold higher binding capacity of the wt. The apparent affinity (i.e. the protein concentration needed to reach half of the maximum protein binding in our experimental setup) is comparable for M1 and M1-N, but significantly higher for M1_m_. Altogether, SPR measurements indicate that the N-terminal domain of M1 has an affinity to DOPS-containing membranes similar to that of the whole M1 protein. Modification of the PBD (as in M1_m_) significantly decreases protein binding to the membrane. Finally, no binding could be measured between M1-C and DOPS-containing monolayers at protein concentrations up to ca. 130 μM. More specifically, the binding of M1-C to DOPC/DOPS monolayers was in general very low (<500 RU at 30 μM, compared to ca. 10000 RU for M1 at 1 μM) and comparable to that of the reference surface control containing the positively charged DOTAP lipid.

**Table 1.** Parameters obtained from SPR binding curves of the different M1 constructs. Binding curves as represented for example in Fig. 2B were analyzed using equation (1). *R_∞_* is the amount of RU at the maximum coverage of bound protein, and 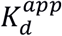 is an apparent dissociation constant of the protein to the lipid monolayer, i.e. the protein concentration at which half of the accessible monolayer surface is occupied. *n* represents the number of independent protein preparations that were analyzed on different days. For each protein, we calculated the average and the standard deviation of each parameter.

| Construct | $R_{\infty}$ (RU) | $K_d^{app}$ (nM) | $n$ |
| --- | --- | --- | --- |
| M1-wt | 8000±3000 | 100±20 | 5 |
| M1<sub>m</sub> | 2900±900 | 290±80 | 4 |
| M1-N | 7200±400 | 70±20 | 4 |
| M1-C | -180±80 | N/A | 3 |

### Conformation of full-length M1 and its C-terminal domain is modulated by membrane binding

To compare the structural rearrangement of M1 constructs upon membrane interaction, CD measurements were performed in the presence and absence of liposomes. In this experimental setup, large unilamellar vesicles (LUVs) of different lipid compositions were used to study M1 membrane binding. DOPS-containing LUVs were compared to neutral liposomes prepared from pure DOPC. For all four protein constructs (i.e. M1, M1_m_, M1-N and M1-C), CD spectra obtained in the presence of DOPC-liposomes did not differ from CD spectra obtained in buffer (Fig. 3), indicating that there either are no interactions with neutral liposomes or that any such interactions are not accompanied by changes in secondary structure composition. These results are consistent with previously reported CD spectra of M1 constructs in solution [33, 37, 47] and with former findings indicating only very weak association of M1 with neutral PC membranes [21, 32]. The estimated secondary structure contents were found to be comparable to those previously reported [33] (data not shown). In the presence of negatively charged liposomes containing DOPS, however, the spectra of three of the constructs (i.e. M1, M1_m_ and M1-C) displayed moderate, yet significant alterations compared to measurements performed in the absence of lipid vesicles. Such alterations might be brought about either by small structural rearrangements in a large number of molecules that associate with the liposomes or by limited binding of a subpopulation of protein molecules that is accompanied by more pronounced structural rearrangements. The CD data, therefore, do not provide direct information about the membrane affinities of the different constructs (i.e. the fractions of bound protein). For the N-terminal domain alone (M1-N), no conformational change could be observed in the presence of DOPS-containing lipid bilayers. This does not necessarily exclude binding of M1-N to these membranes, but rather suggests that—if there is binding, as observed in SPR measurements (Fig. 2) —this is not accompanied by modifications of secondary structure.

### Both M1 N-terminal and C-terminal domains are needed for efficient protein multimerization upon interaction with lipid membranes

Fluorescence laser scanning microscopy can also be used to measure protein–protein interaction for the different fluorescent M1 constructs bound to SLBs. We performed RICS analysis, a quantitative fluorescence microscopy approach that provides diffusion coefficients and the relative molecular brightness of membrane-bound proteins assemblies [32]. The latter is a measure of the multimerization state of the protein. To this end, stacks of 100 images were acquired, and independently moving fluorescent entities were detected by analysing the fluorescence fluctuations among pixels. Note that these independently diffusing objects can be monomers or multimeric complexes, which yield different brightness values. The brightness is defined as the total fluorescence intensity of membrane-bound protein divided by the number of protein clusters detected, thus indicating the degree of clustering in the sample (or the relative amount of M1 molecules per cluster). On the one hand, higher-order multimers are characterized by higher brightness values. On the other hand, the diffusion coefficient provides information regarding both size of protein assemblies (i.e. large protein multimers diffuse slowly) and protein-membrane interaction (i.e. proteins loosely bound to the membrane surface will diffuse faster [48]).

As shown in Fig. 4A, M1 wt bound to SLBs containing 30 mol% DOPS has a diffusion coefficient of (0.29 ± 0.03) μm^2^/s, in agreement with previous estimates (32), while the diffusion coefficient of M1-C was found to be significantly larger at (0.91 ± 0.07) μm^2^/s, clearly indicating slower dynamics of wt M1, as compared to M1-C. Consistently, the brightness of wt M1 was ~4.3-fold higher than that of M1-C, suggesting that M1-C has a lower tendency to oligomerize as compared to full-length M1 (Fig. 4B). Diffusion coefficients of M1_m_ and M1-N were found to be closer to wt values (only ~2-fold higher in average). Also, the brightness of M1_m_ was not significantly different from the brightness of full-length M1 wt, suggesting oligomerization of both constructs to be similar.

**FIGURE 4.**
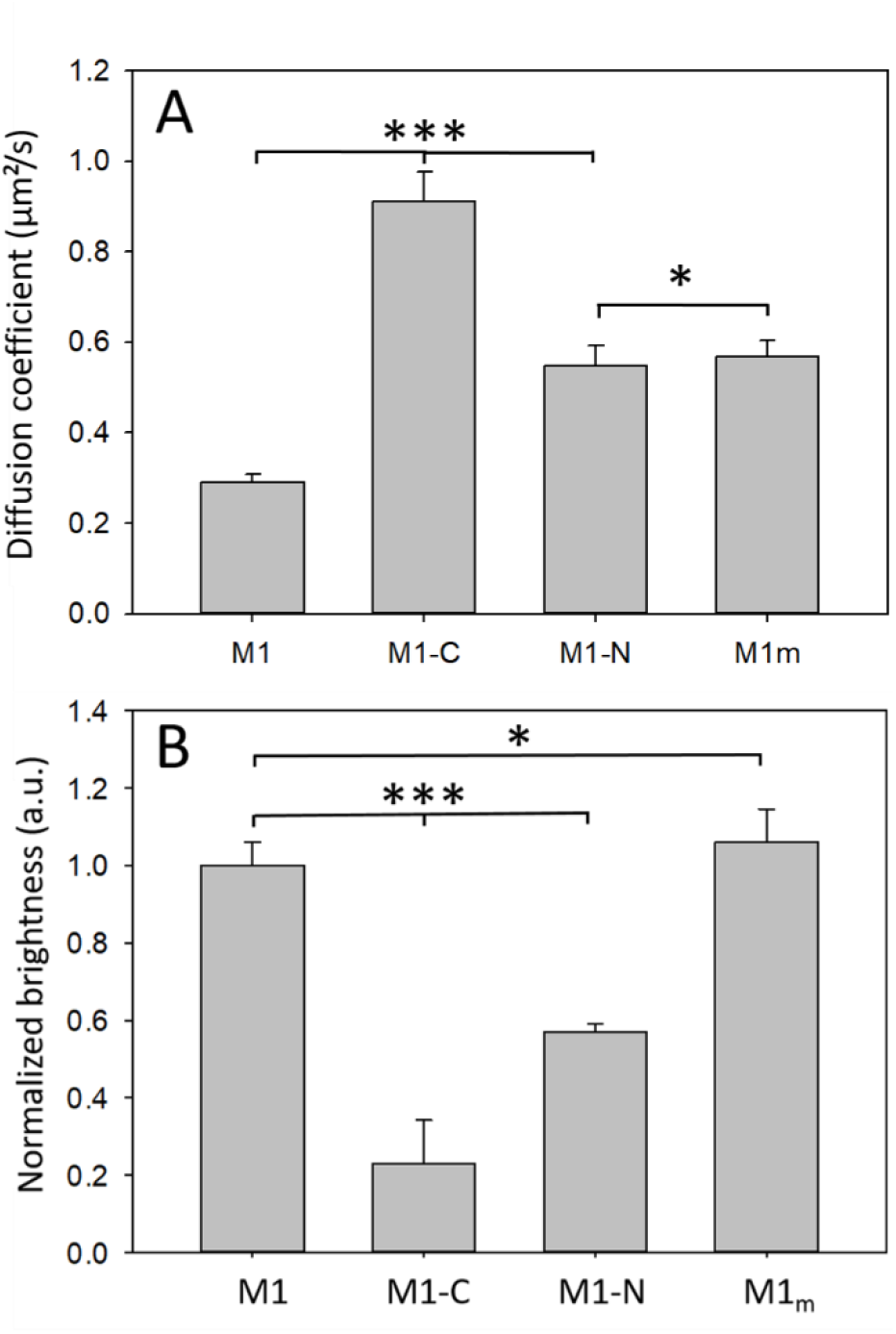
RICS analysis of diffusion and multimerization of fluorescently labeled M1 constructs. Images of M1 constructs (50 nM) bound to SBLs consisting of 70 mol% DOPC and 30 mol% DOPS were analyzed via RICS. A: Diffusion coefficients of protein constructs M1 wt, M1-C, M1-N and M1_m_ on the membrane surface. B: Normalized fluorescence brightness was determined from the same samples as in panel A. Data were normalized to brightness values of M1 wt. Error bars represent the standard error of the mean (SEM) (*n*≈100-200 distinct image stacks from 3 (M1, M1-N, M1m) or 4 (M1-C) sample preparations). Asterisks indicate statistically significant differences determined with a two-sided t-test with distinct variances: (^***^ corresponds to p<0.01; ^**^ corresponds to p<0.05; ^*^ corresponds to p>0.01, i.e. difference is not statistically significant). All measurements were conducted at room temperature.

### Structure and intramolecular dynamics of full-length M1 monomer obtained via MDS

As a complementation to our experimental investigations, we performed MDS in order to gain information on the details of M1–lipid interaction at the atomic level. We first started with the analysis of a single M1 monomer in solution. One possible model for the structure of the full-length protein (Fig. 5A and B) was obtained by combining the known structure of M1 N-terminal domain (PDB ID 1EA3), composed of α-helices H1 to H10, with separate *ab initio* modelling of the C-terminal domain. Furthermore, we calculated the minimum free-energy orientation of the C-terminal domain with respect to the N-terminal domain, as described in the Materials and Methods section. The structure obtained for M1-C consists of three α-helices (namely, H11, H12 and H13), in agreement with a previous prediction based on the M1-C amino acid sequence and bioinformatics tools [37]. In our model, the three M1-C α-helices align approximatively parallel to the M1-N surface provided by α-helices H1, H2, H7 and H9, in line with previous reports [49].

**FIGURE 5.**
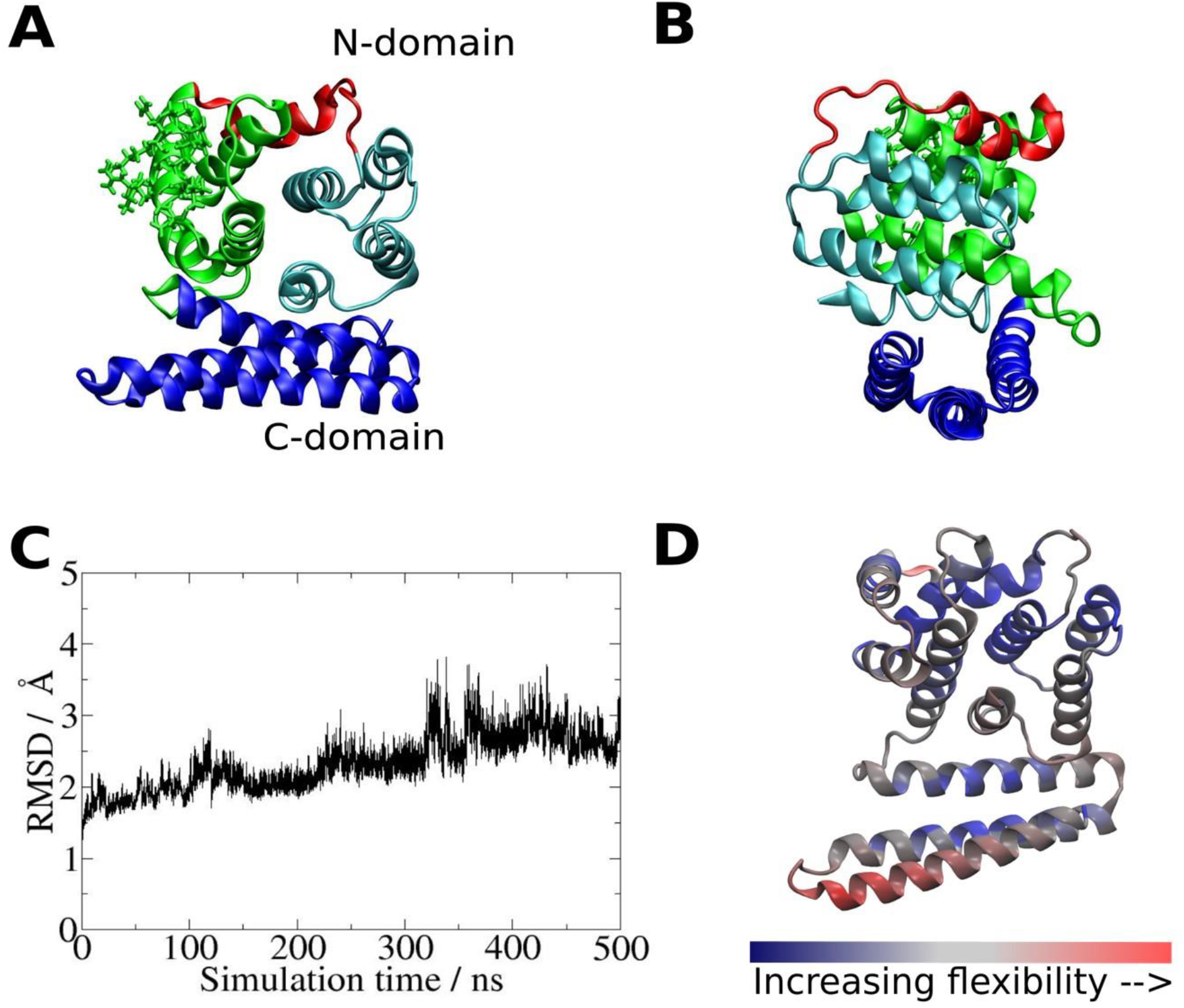
Model of M1 structure and molecular dynamics simulation (MDS) of full-length M1 in solution. A: Model of the M1 monomer in solution. The proposed structure of the M1-C domain (blue, separately obtained by *ab initio* modelling) in association with the crystal structure of the M1-N domain (green, red, cyan; PDB 1EA3) obtained after a 500 ns MDS run is presented. The M1-C domain consists of three α-helices, namely, H11, H12 and H13. Basic residues in the polybasic domain (PBD, aa 95–105) are displayed in stick representation. B: Same model as in panel A, but rotated by 90°. C: Root-mean-squared deviation (RMSD) values for the whole M1 protein simulation time scale. D: Representation of the M1 structure with the corresponding flexibilities of each region as obtained by normal-mode analysis. Colours represent the mode amplitude (i.e. flexibility) for the first normal mode (blue: low flexibility; red: high flexibility).

In order to test the stability and dynamics of the M1 structure, a 500-ns MDS of the full-length protein was performed. The protein reached a stable conformation after ca. 400 ns, as shown by the root-mean-squared displacement (RMSD) values depicted in Fig. 5C. Moreover, normal-mode analysis [50] of residue dynamics revealed a “hot”, highly flexible region among the helices in the M1-C domain characterized by a large amplitude of the first “breathing” mode (highlighted red in Fig. 5D), while the other regions of the protein remained comparatively rigid (highlighted in blue). The observed flexibility of the C-terminal domain is in agreement with a previous theoretical analysis [37] and small-angle X-ray scattering investigations [36]. Another small, but flexible loop can be further observed between helices H5 and H6 of the N-terminal domain (aa 82–86, Fig. 5D).

### Binding of an M1 monomer to a model membrane studied by MDS

To obtain more precise information regarding the binding of a single M1 monomer to a lipid membrane, various spatial configurations of the M1–membrane system were evaluated, as described in the Materials and Methods section and Fig. S2. Briefly, the protein was slowly translated towards the membrane surface using eight different orientations relative to the membrane plane. Upon reaching the membrane, only one orientation resulted in a non-diverging MDS and a stable interaction between the protein and the DOPC/DOPS model membrane. The other seven orientations tested led to configurations in which the protein did not remain bound to the membrane (in the absence of a pulling force), as shown in Fig. S3. The protein–membrane system exhibiting the most stable interaction (the second in Fig. S2) was further simulated for a total of 500 ns. The final result shown in Fig. 6A and B indicates that the M1 N-terminal domain mediates the interaction with the membrane, while C-terminal domain does not directly interact with lipids (i.e., it is on the distal side of the protein). More specifically, in our model, the membrane-interacting region was found to include helix H5 (aa 78 to 84) and two surrounding loops (aa 68 to 78 and 84 to 88) that approximately correspond to the region denoted in red in Fig. 6A and B. Arginine and glutamine residues from this region establish direct interactions with components of the membrane. In detail, arginine residues interact with PS head groups, while two glutamine residues (Gln75 and Gln81) display interactions with PC head groups. Some of these interactions can be seen in closer detail in Fig. 6C. As shown in Fig. 6D, the distances between the glutamine residues and the closest PC head groups (N–P atom distances) consistently reach values as low as 4 Å, indicative of transient but close interactions [51].

**FIGURE 6.**
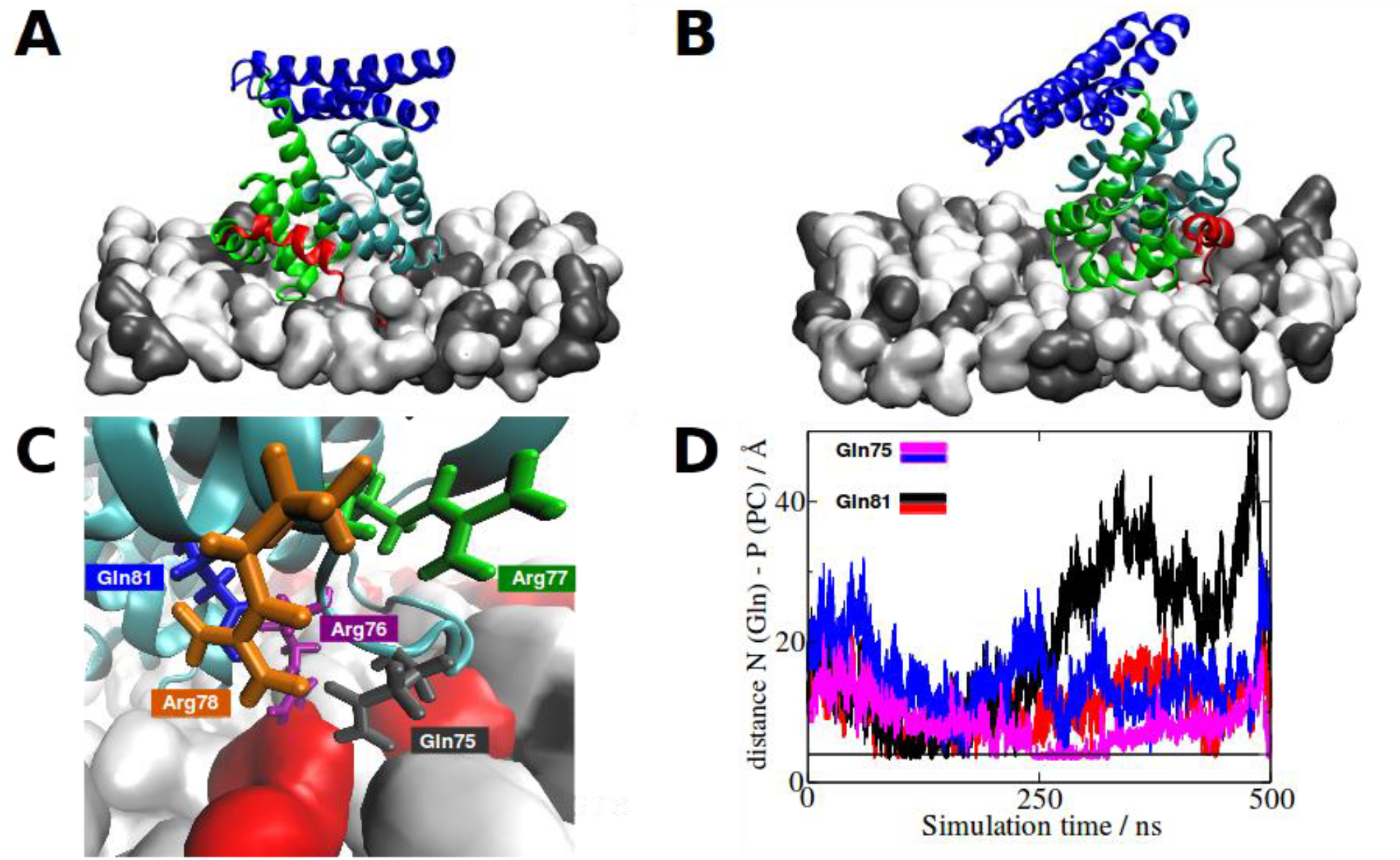
Binding of M1 to a lipid bilayer investigated by MDS. The most stable orientation of M1 binding to a DOPC/DOPS (70 mol%/30 mol%) bilayer is presented. A: Representation of the lowest free-energy configuration of a M1 monomer bound to a model membrane. Colour code corresponds to that of Fig. 5A; head group of DOPC (white), phosphoserine head group of DOPS (dark grey). For simplification, water molecules, ions, lipid tails and the opposite monolayer of the lipid bilayer are not shown. B: 90°-rotated representation of A. C: Closer view of the interactions between Arg78 and a phosphoserine head group of DOPS, and of Gln81 with a phosphocholine head group of DOPC. For a better visualization in this panel, PS head groups are evidenced in red. D: Distance values over simulation time scale between either Gln75 or Gln81 and the phosphate group of the two nearest PC head groups. The distances between the N in the Gln side chain and the P in the PC head are represented as a function of time. The horizontal line corresponds to 4 Å.

Interestingly, the PBD of M1 on helix 6 (displayed in green in Fig. 5A, with basic residues in stick representation) is not stably involved in direct interactions between the protein and the membrane in our model. More intriguingly, another region on helix 5 including the arginine triplet in position 76-77-78 appears to be interacting specifically with the negatively charged head group of DOPS during the whole simulation (Fig. 6C). The positively charged side chains of Arg76 and Arg78 are consistently involved in a short-distance interaction with the head groups of DOPS molecules. Distances between arginine residues 76, 77 and 78 and the three closest PS head groups are shown in Fig. 7A–C. Arg76 and Arg78—but not Arg77—are found in close proximity to a PS head group during several periods of the simulation (e.g., green curves in panels A and C, often and consistently below 3–4 Å), suggesting strong protein–lipid interactions [52]. In order to better quantify these interactions, we calculated radial distribution functions, as previously described [53]. The curves shown in Fig. 7D indicate a high probability to find a P atom of a DOPS molecule at short distances from Arg 76 and 78 (e.g., probability peak at 2.7 Å distance). The same analysis for Arg77 does not suggest a specific interaction with DOPS molecules. Similar interactions among adjacent arginine residues and acidic epitopes have been also previously described [54].

**FIGURE 7.**
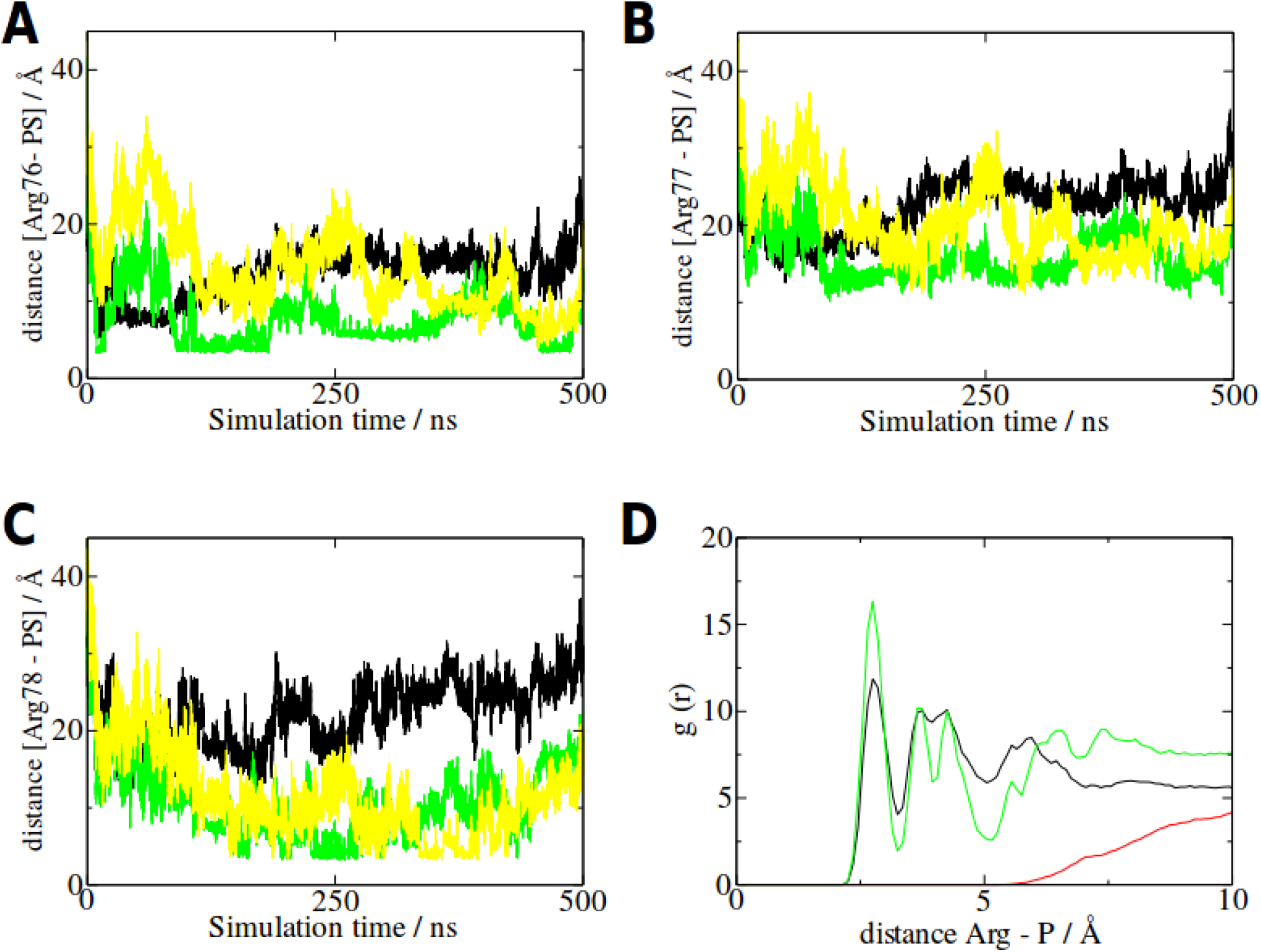
Interaction between M1 76-78 arginine residues and closest PS lipid head groups during the simulation time scale. A–C: Distances between arginine (Carbon Z) 76, 77 and 78, and the P atom of the three closest PS head groups, as a function of simulation time. Each colour represents a different PS molecule in the membrane. D: Radial distribution probability functions for the distance between the centre of mass of the Arg76 (black), Arg77 (red) and Arg78 (green) and the phosphor atom (P) of DOPS in the membrane layer. While the curves corresponding to Arg76 and 78 present a first large peak around 2.7 Å, indicating a high possibility of finding a phosphorus atom of a DOPS during the simulation time scale, no such pattern is observed for Arg77.

### Binding to lipid membranes restricts M1 intramolecular dynamics

A comparison of residue dynamics of M1 in solution and in the membrane-bound state can be performed by comparing the root-mean-squared fluctuations (RMSF) calculated from equilibrium simulations for both systems, as presented in Fig. 8. In agreement with the results obtained from normal mode analysis (Fig. 5D), the most flexible region of M1 in solution was found in the C-terminal domain of the protein, featuring an RMSF peak greater than 3 Å. Interestingly, an overall reduction of the fluctuation was observed for the full-length protein upon interaction with the lipid membrane (blue curve in Fig. 8). Notwithstanding the fact that the association of M1 with the membrane occurs through the N-terminal domain, binding of the protein to lipids seems to induce a significant stabilization also of the C-terminal domain, as noted by the reduction in the flexibility observed for this region. Notably, the peak corresponding to residues 82–86 of the N-terminus, which in the protein structure represents the loop neighbouring the membrane-bound triplet of arginine residues and membrane-interacting glutamine residues (Fig. 6), is significantly reduced upon membrane association of the protein.

**FIGURE 8.**
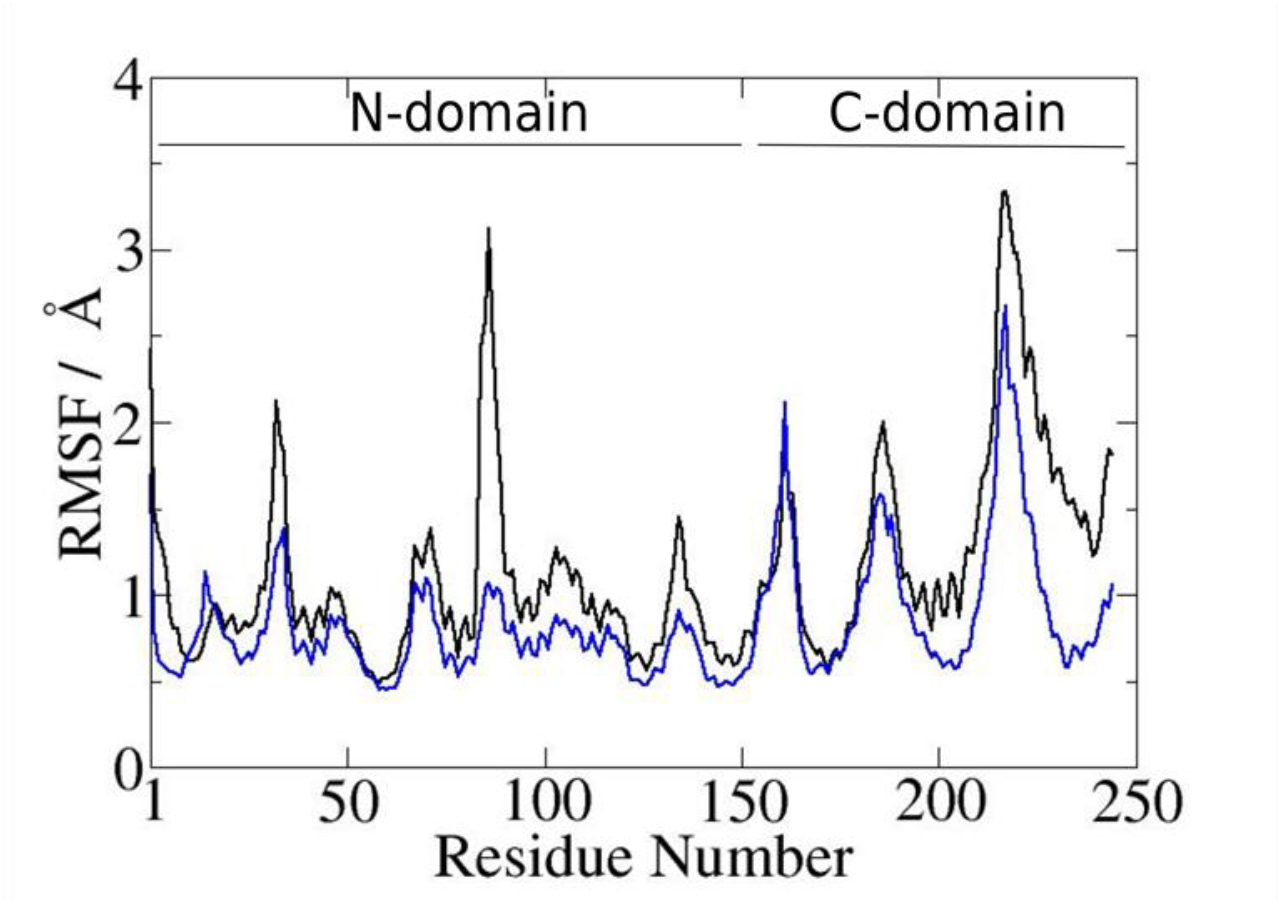
RMSF of M1 residues are reduced upon binding to membrane. RMSF is plotted as a function of amino acid sequence in M1 as calculated for the protein in free solution (black) or bound to the model membrane (blue). Values correspond to mean values along the 500-ns simulation time scale.

## DISCUSSION

The assembly of IAV at the PM of infected cells is a complex process governed by multiple and inter-dependent protein–protein and protein–lipid interactions. Specifically, M1 is supposed to orchestrate the assembly process by concurrent interactions with viral proteins, vRNPs and lipids in the PM. We and others [5, 21, 29, 32] have characterized the binding between M1 and PS lipids, both in model and cellular systems. More in detail, we have recently proposed a model according to which M1-M1 interactions (i.e. protein multimerization) is enhanced by M1–lipid binding, possibly via alterations in protein structure. While M1 multimerization in the absence of lipid membranes has been found to occur at relatively high concentrations (ca. 30 μM) and to be regulated by the N-terminal domain of the protein [47], much less information is available regarding the multimerization of lipid-bound M1.

In order to identify the regions of M1 that are responsible for protein–lipid interaction and protein-multimerization in the membrane-bound state, we produced three M1-derived constructs: M1-N (N-terminal fragment, aa 1-164), M1-C (i.e. C-terminal fragment, aa 165-254) and M1_m_ (in which the basic residues in the PBD were mutated to alanine residues, 95-AAVALYAALAA-105). Using a similar approach, a qualitative characterization of M1–lipid interaction has previously suggested that the N-terminal domain plays a fundamental role [21], although no consensus was reached regarding the specific regions actually mediating M1–membrane binding [28, 55, 56]. Here, we quantified the binding of the M1 constructs and full-length M1 wt to PS-containing model membranes using a combination of SPR and fluorescence microscopy. The advantage of this combined approach is that fluorescence-based analysis can provide, at the same time, distinct data regarding protein–protein and protein– lipid interactions. RICS allows in fact a quantification of total protein binding to the SLB (i.e., by normalized fluorescence intensity, Fig. 1), formation of multimers (i.e., by increase in normalized molecular brightness, Fig. 4B) and protein dynamics (i.e., by diffusion coefficients of M1 multimers, Fig. 4A). Conversely, SPR quantification of protein binding does not require labeling of M1, but cannot straightforwardly distinguish binding of monomers or small oligomers and the formation of larger multimers at the lipid interface, since only the total amount of bound protein is measured [57].

### Interaction between M1-C and PS-containing lipid membranes

Both SPR and fluorescence experiments show that M1-C binds to PS-containing membranes with much lower affinity than M1 wt. Fig. 1 shows a ~90% lower binding compared to M1 wt. Similarly, no significant binding of M1-C to PS-rich lipid monolayers could be detected via SPR. The observation that the binding of the C-terminal domain of M1 is at least one order of magnitude lower than that of the full-length protein is qualitatively confirmed also by earlier investigations [21]. Our RICS data also show that M1-C bound to the lipid bilayer surface features much faster lateral dynamics and lower molecular brightness (compared to the other protein constructs), thus strongly suggesting that the C-terminal domain of M1 is only loosely bound to the membrane surface and is not capable of forming large multimers. Interestingly, although the interaction with lipid membranes is quite weak, CD spectra (Fig. 3) in the presence of PS-containing liposomes are different from those recorded in the absence of lipid membranes. This observation suggests a conformational adaption in the spatial configuration of M1-C upon interaction with lipids, although a more quantitative characterization cannot be provided by the available data. Due to the limited degree of multimerization, it is safe to assume that the structural alterations detected by CD in the case of M1-C samples are mostly due to protein–lipid interaction, rather than protein–protein interaction.

### Interaction between M1_m_ and PS-containing lipid membranes

Similarly to the case of M1-C, both fluorescence microscopy and SPR report a weak interaction between M1_m_ and PS-containing lipid membranes, compared to M1 wt. Fluorescence intensity quantification (Fig. 1) indicates an 80% decrease of protein binding at 50 nM concentration compared to M1 wt. Likewise, SPR data (Fig. 2 and Table 1) show that the apparent *K_d_* of M1_m_ is ca. 3–4 times higher than that of M1 wt. The low response measured by SPR cannot be ascribed, for example, to limited multimerization and protein– protein interaction of M1_m_ compared to M1 wt, since RICS data suggest that the two proteins multimerize to a similar extent (Fig. 4B). The observed differences must therefore be connected to a less stable interaction between M1_m_ and PS-rich membranes, as confirmed by a significantly higher diffusion coefficient compared to that of M1 wt (Fig. 4A). Taken together, these data suggest that membrane-bound protein multimers formed by M1_m_ are fewer but of similar size to those formed by M1 wt.

The PBD, which is modified in M1_m_, has been extensively studied in the past two decades and has been shown to be essential for virus replication [11, 58, 59]. The ability of this amino acid stretch to interact with negatively charged lipid membranes was suggested on the basis of liposome-binding experiments [21], although its role in mediating membrane binding in a cellular context remains controversial. A number of studies, including mutational analyses in a cellular context and tritium bombardment of intact virus particles, suggest that M1 membrane association is mediated not only by the PBD, but rather by cooperative action of several binding sites [60, 61]. Our results support this conclusion, suggesting that the positively charged residues in the PBD are involved (although not strictly necessary) in M1– lipid interactions. Finally, CD data indicate that structural rearrangements of M1_m_ occurring concomitantly to membrane binding are very similar to those measured for the wt protein (discussed below) and, therefore, do not seem to be influenced by the PBD.

### Interactions between M1-N and PS-containing lipid membranes

Fluorescence-based quantification of M1-N binding to PS-containing lipid membranes indicates that this construct binds only to limited extent (i.e., ~20% compared to the full-length protein). On the other hand, SPR data (Fig. 2 and Table 1) clearly indicate that M1-N and M1 wt have similar K_d_ values (i.e., ~80−90 nM). One of the reasons for such discrepancy might be the presence of the fluorescent label covalently bound to randomly exposed primary amines in the M1-N construct which was used for fluorescence measurements. Due to the shorter length of the construct, it is more probable that the fluorescent label binds to those residues that are determinant for membrane binding. In line with this interpretation, our MDS data (Fig. 6C) show that interactions between lipids and full-length M1 might involve residues belonging to N-terminal domain (and therefore to M1-N as well) that possess aliphatic amines able to react with the succinimidyl ester group in the fluorescent label, such as Gln75, Gln81, Arg76 and Arg77. Notably, a recent SPR study has shown that labeling of wheat germ agglutinin with fluorescent groups reduces significantly binding affinity of this lectin [62]. Unfortunately, we could not study binding of labeled M1-N via SPR since, due to the rather low labeling efficiency (see above), SPR would report essentially on the non-labeled protein fraction. Efforts to increase labeling failed as protein aggregation became an issue. Thus, in this specific case, we argue that label-free SPR quantification is more reliable when quantifying M1-N binding to PS-containing membranes. RICS characterization of protein dynamics and multimerization state (Fig. 4) reports values that are intermediate between M1 wt and M1-C. Considering once more a possible interference caused by fluorescence labeling (and therefore an underestimation of the multimerization state), we can nevertheless conclude that M1-N is capable of forming multimers to a degree similar to or slightly lower than the full-length M1. This conclusion is fully supported by previous observations on virion structure, showing that the N-terminus of M1 is the main determinant of M1 multimerization, although the C-terminal domain makes also a significant contribution [22]. Finally, CD spectroscopy (Fig. 3) indicates that binding of M1-N to lipid membranes is not accompanied by major changes in secondary structure.

### Full-length M1 and its interactions with PS-containing lipid membranes

The interactions between M1 wt and PS-containing lipid membranes have recently been characterized using RICS [32]. In the current work, we could confirm the efficient binding of M1 to negatively charged membranes and the significant formation of multimers at 1–100 nM concentrations (Figs. 1 and 4). Here, we report additional information regarding the structure of the full-length protein, both in solution and as a monomer bound to a lipid bilayer. While the M1 N-terminus has been crystallized and described as globular and ordered [33, 63], no structural data are currently available regarding the C-terminal domain. Studies involving CD, bioinformatics and tritium labeling have described the C-terminal domain as being composed of three α-helices and characterized by a high level of structural disorder [37]. Small-angle neutron scattering [33] and a recent low-resolution reconstruction of the full-length M1 from small-angle X-ray scattering suggest that the protein has an elongated shape with a globular N-terminal domain and an elongated and flexible C-terminal domain [36]. Finally, very recent small-angle X-ray scattering (including ROSETTA modelling and docking analysis) confirmed these general findings and proposed the existence of multiple structural configurations that the flexible C-terminal domain can adopt [64].

In this work, we further address the lack of high-resolution information regarding the structure of full-length M1 by means of MDS. Our data confirm that the C-terminal domain consists of three α-helices, and we further characterized their orientation with respect to the N-terminal domain (Fig. 5A and B). Quantification of the RSMF and normal mode analysis suggest, in line with previous results [36], a significant flexibility of the C-terminal domain, most notable in the region between helices H12 and H13 (Figs. 5D and 8). A remarkable flexibility is further suggested also for a loop between H5 and H6 of the N-terminus. It is worth noting that the M1 structure described here is to be considered as one of several possible conformations that the protein can adopt. This assumption is clearly corroborated by the similarity between the M1 structural model presented here and one of several possible M1 structures that were recently described [64]. Our model of M1 in solution was then used to analyze M1–membrane interaction by MDS and to provide an interpretation of our experimental data on a molecular and atomistic level.

The interaction between a single M1 molecule and a PS-containing lipid membrane was thus analyzed via MDS. Our data indicate that M1–lipid binding might occur through the N-terminal domain and, in particular, through the region around H5. Here, we could identify stable interactions between PS and Arg76/Arg78 and between PC and Gln75/Gln81 (Figs. 6 and 7). These results are in full agreement with our experimental characterization of the membrane-binding capabilities of the isolated M1 N-terminal (i.e., significant membrane binding) or C-terminal (i.e., no detectable binding) domains. Also, tritium planigraphy studies of M1 in virions [65] located the contact surface between protein and viral membrane approximatively in the same region of the protein (i.e., around aa 90). Furthermore, a recent study indicated that Arg75–78 are essential for M1 binding to model lipid and cellular membranes [66]. Nevertheless, it should be noted that the specific orientation between a single M1 and the lipid membrane that we report here is only one of the most stable ones according to our calculations: other spatial configurations might also be possible, especially in the presence of concurrent protein–protein and protein–lipid interactions. For example, our experimental data on M1_m_ and previous findings indicate that other regions of the protein (e.g. PBD) might act synergistically in mediating M1–lipid interaction, as such versatility is needed for M1 to associate with different cellular organelles and compartments during infection [60].

Following the binding to the lipid membrane, the flexibility (i.e. the spatial fluctuations) of the C-terminal domain is significantly reduced, as suggested by MDS (Fig. 7). This is particularly surprising, since protein–lipid interaction occurs through the N-terminal domain, whereas the C-terminal domain is not directly involved. The modulation of intramolecular chain dynamics might therefore be brought about by a re-organization of the interaction network among amino acid residues, resulting from the binding to the lipid bilayer. In this context, N-terminal residues 82–86, which also show a remarkable loss in flexibility upon membrane binding, might play a role (Fig. 8). This site is situated between the membrane-interacting PDB, aa 95–105, (Figs. 1 and 2) and the membrane-interacting arginine- and glutamine-rich sequence, aa 75–81 (Fig. 6). Also, our CD data suggest that moderate but significant structural modifications occur upon binding of M1 wt to lipid membranes (Fig. 3). Such modifications are not observed for M1-N alone and, therefore, might involve specifically the full-length protein or the C-terminal domain. The fact that no significant structural rearrangements are observed for a single membrane-bound M1 molecule by MDS, suggests that the changes observed in the CD spectra of full-length M1 might be caused by protein–protein interactions, occurring concurrently to protein–lipid interactions. The structural plasticity of M1 as previously discussed [36, 37, 64] and, in particular, conformational changes in the full-length protein as we report here (especially in the C-terminal domain) might be required for the multi-functionality of M1, including its ability to multimerize and mediate diverse steps in the infection process.

## CONCLUSIONS

We have applied a combination of fluorescence microscopy, SPR, CD and MDS to characterize M1–M1 and M1–lipid interactions from a structural point of view. Our results indicate that binding of M1 to lipid membranes occurs mainly through interactions between specific residues of the N-terminus (e.g., Arg76–78) and polar lipid head groups. Also, it is suggested that the conformational flexibility of the M1 C-terminal domain decreases as a consequence of binding to the lipid membrane. Furthermore, protein–protein and protein– lipid interactions occurring during membrane binding of M1 are accompanied by a moderate but significant alteration in secondary structure. These findings support a model of M1–M1 interactions that we and others have discussed [21, 32, 38], according to which binding of M1 to the PM (mediated by the N-terminal domain) could affect the structure or the chain dynamics of the protein (specifically of the C-terminal domain), thus exposing or stabilizing the interfaces for M1 self-association [32] and, possibly, vRNP binding [21].

## CONTRIBUTIONS

C.T.H., S.D.L., N.J., S.K., A.H. and S.C. designed the experimental plan. C.T.H., S.D.L., I.D., N.J., N.B., S.B. and Q.H. performed experiments and simulations. C.T.H., S.D.L., I.D., N.J., N.B., S.B., Q.H. and S.C. analyzed the data. C.T.H., S.D.L., I.D., Q.H., S.K., A.H. and S.C. wrote the manuscript.

## ACKNOWLEDGMENTS

The authors thank Svenja Nierwetberg for help with protein purification. This work was supported by the German Research Foundation (DFG) (grants CH 1238/3 to S.C., KE 1478/4 to S. K and SFB 765 to A.H.) and the Natural Science Foundation of China (grant 91430112 to Q.H.). S.D.L. is a Georg Forster Fellow of the Alexander-von-Humboldt Foundation in Germany.

